# Fixational drift is driven by diffusive dynamics in central neural circuitry

**DOI:** 10.1101/2021.02.10.430643

**Authors:** Nadav Ben-Shushan, Nimrod Shaham, Mati Joshua, Yoram Burak

## Abstract

During fixation and between saccades, our eyes undergo diffusive random motion called fixational drift [1]. The role of fixational drift in visual coding and inference has been debated in the past few decades, but the mechanisms that underlie this motion remained unknown. In particular, it has been unclear whether fixational drift arises from peripheral sources, or from central sources within the brain. Here we show that fixational drift is correlated with neural activity, and identify its origin in central neural circuitry within the oculomotor system. We analyzed a large data set of ocular motoneuron (OMN) recordings in the rhesus monkey, alongside precise measurements of eye position [2, 3], and found that most of the variance of fixational eye drifts must arise upstream of the OMNs. The diffusive statistics of the motion points to the oculomotor integrator, a memory circuit responsible for holding the eyes still between saccades, as a likely source of the motion. Theoretical modeling, constrained by the parameters of the primate oculomotor system, supports this hypothesis by accounting for the amplitude as well as the statistics of the motion. Thus, we propose that fixational ocular drift provides a direct observation of diffusive dynamics in a neural circuit responsible for storage of continuous parameter memory in persistent neural activity. The identification of a mechanistic origin for fixational drift is likely to advance the understanding of its role in visual processing and inference.

## Main

In order to explore the fine details of a visual scene, we fixate our gaze on specific areas of interest [4]. Yet even during fixation the eyes are not completely stationary. Over intervals that typically last a few hundred milliseconds, the eyes exhibit continuous motion called fixational drift, flanked by microsaccades [1, 5]. Eye trajectories during fixational drift are smooth, but are highly variable and are characterized by the statistics of a super-diffusive random walk [6, 7, 8, 9]. The role of this irregular smooth motion in vision has been extensively studied and debated in the past few decades. It has been proposed that fixational drift increases the information carried by retinal spikes on the visual stimulus, thereby aiding high acuity vision [5, 10, 11, 12, 13, 14]. On the other hand, it has been argued that fixational drift poses a computational challenge for high acuity inference in the visual cortex, since attempting to overcome retinal spiking noise by simple temporal averaging would smear out fine visual features [13, 15, 16].

While extensive effort has been devoted to understand the functional role of fixational drift in vision, the mechanisms responsible for this motion have remained unidentified. It is not even known whether the origin of fixational drift is peripheral – arising, e.g., from noisy dynamics of the ocular muscles [17], or whether fixational drift arises in more central brain circuits in similarity to saccades [1, 7, 18]. The main difficulty arises from the small amplitude of fixational drift in comparison with other types of eye movement. In human subjects it is highly challenging to measure the minute details of this motion [4], and measurements of single neuron activity are not available. Therefore, we focus from here on on non-human primates, whose oculomotor system is highly similar to that of humans [4].

During smooth pursuit and saccades, eye trajectories can be predicted quite precisely from the spiking activity of single oculomotor neurons (OMNs). Correspondence between eye movements and neural activity has been established also during microssacades [19]. However, a correspondence between the position of the eye and single OMN activity has not been demonstrated during fixational drift. It is highly challenging to test for such a relationship, because the subtle changes that are expected to occur in the firing rate of individual OMNs during fixational drift are largely masked by their spiking noise.

Here we report that a systematic relationship does exist between fixational eye motion and OMN activity. Moreover, most of the variability in fixational eye drifts arises in central neural circuitry upstream of the OMNs. Using a theoretical model, constrained by the parameters of the primate oculomotor system, we point to a likely source of this motion in the oculomotor integrator, a memory circuit in the brainstem which is responsible for maintenance of a steady eye position between saccades. Thus, we propose that fixational eye drifts are driven by random diffusion along a line attractor neural network, as predicted by theoretical works that examined how noise influences the maintenance of continuous-parameter working memory in the activity of neural circuits.

To test for a systematic relationship between OMN activity and fixational eye drifts, we analyzed a large data set of OMN extracelluar recordings in the rhesus monkey. These were collected simultaneously with precise measurements of eye trajectories using a search coil. Recordings were made while two monkeys moved their eyes to track repeated presentations of a target presented on a screen, initially at rest and then moving at constant speed (Fig.1a). First, we fitted a linear combination of eye position, velocity and acceleration to predict the firing rate of single OMNs during large eye movements [20] (n=57 cells, Fig.1b). Fitted parameters were in agreement with previously reported results [20, 21].

**Figure 1:**
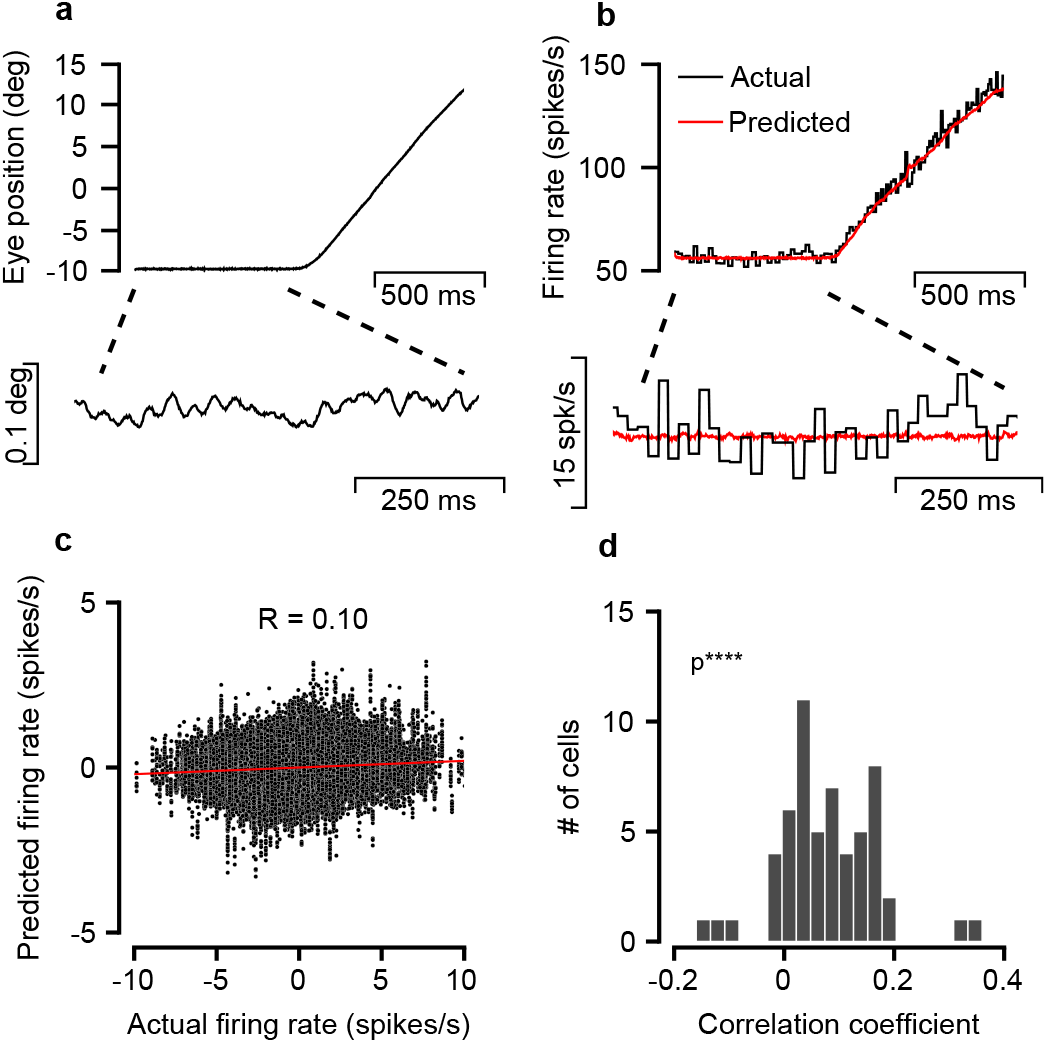
Eye variability during fixation is correlated with motoneuron activity. **a**, Single recorded eye trajectory consisting of fixation followed by smooth pursuit. Lower panel, fixational period expanded and smoothed. **b**, Actual and predicted firing rate of a single OMN during the trajectory shown in (**a**). Lower panel, zoom in on fixational segment. Note that the actual firing rate is much more variable than the predicted rate. **c**, Predicted vs. actual firing rate of a single cell during fixation (mean rate subtracted in both axes). Each point represent the firing rate at a single trial at specific time. Red line: result of linear regression analysis, correlation coefficient: *R=0.10*. **d**, Distribution of correlation coefficients between actual and predicted rate, collected from 57 cells in two monkeys. One sided t-test, *p* = 3 × 10^−9^

OMNs exhibit highly regular firing, with a typical coefficient of variation (CV) of the interspike interval distribution of ~ 0.06 [3]. During fixational drift, however, the variability in the activity of a single OMN is still far too large to identify a correspondence with the small changes in firing rate predicted by eye motion over a single trial (Fig.1b, lower panel). Over the hundreds of trials available from each cell, estimates of the correlation coefficients between the spiking rate of single OMNs and their predictors based on the eye trajectory were noisy (Fig.1c), but significantly deviated from zero over the population of 57 cells (Fig.1d), providing us with initial evidence that neural activity, upstream of the muscles, is correlated with fixational drift.

To facilitate subsequent analysis, we replaced the standard approach discussed above, in which the firing rates of OMNs are predicted from the eye trajectory by a complementary analysis, in which the eye trajectory is predicted from the OMN spikes. This allows us below to quantify shared neural variability directly in terms of its contribution to the eye movements, instead of the firing rates. We thus inverted the fit described above (Methods) to obtain for each OMN a double exponential filter (Fig.2a), whose convolution with the OMN spikes constitutes an estimator of the eye trajectory. Fitted time constants (Fig.2b) were in agreement with previous reports [20, 22]. As expected due to the spiking noise in the activity of individual OMNs, predicted eye trajectories were highly variable compared to the actual eye motion during fixational drift (Fig.2c). Accordingly, correlation coefficients between the predicted and actual eye position differences across 350 ms fixational drift segments (Fig.2d) were small. Furthermore, due to the limited number of trials available for each neuron, the estimates of the correlation coefficients themselves were noisy (gray histogram in Fig.2d). Nevertheless, their average value (〈*R*〉=0.17, one sided *t-test,* p=3 × 10^−9^) was statistically different from zero (Fig.2d, see also Extended Data Fig.1). Overall, our results (Figs.1 and 2) indicated that fixational drift is correlated with neural activity.

**Figure 2:**
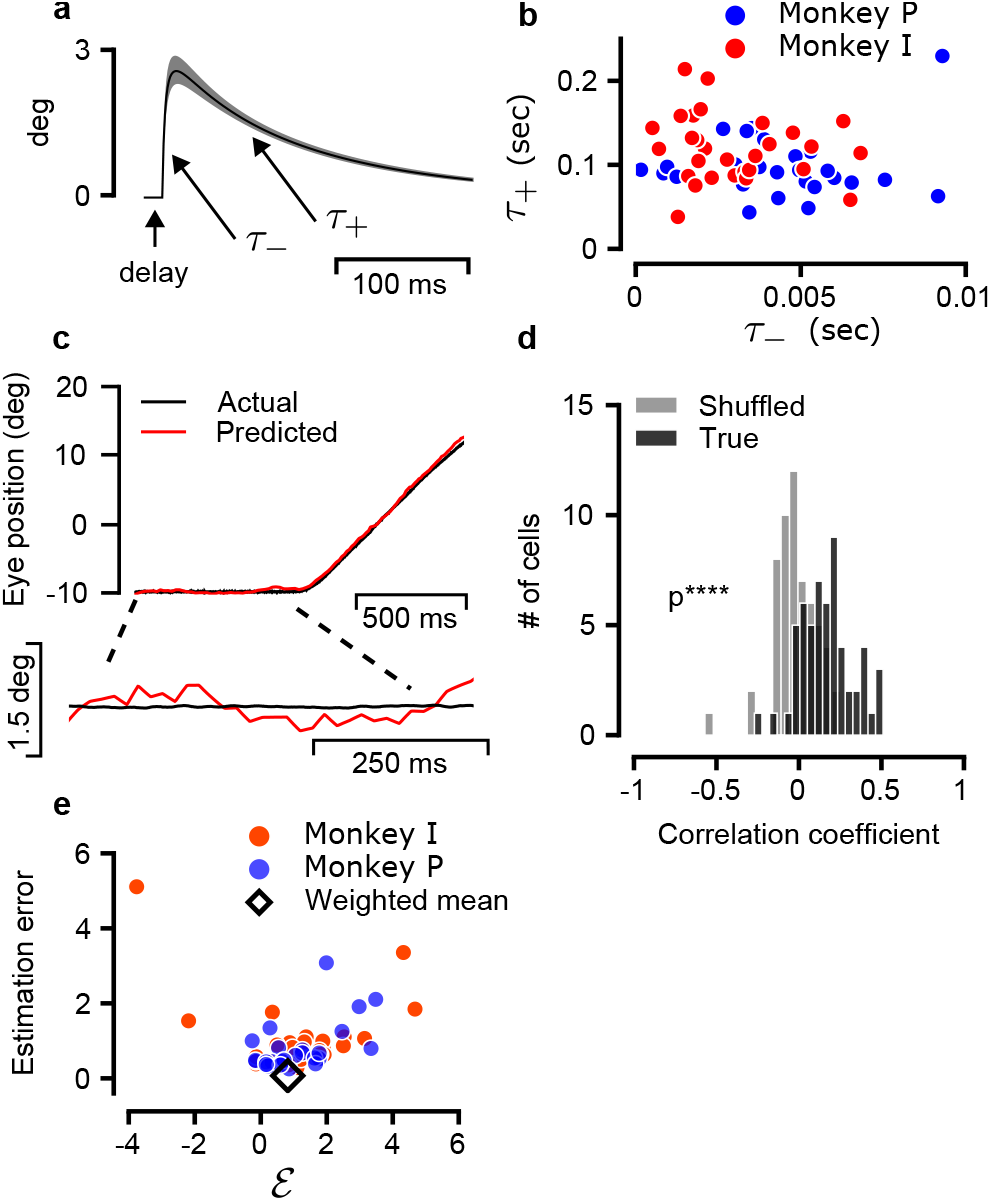
Predicting the eye trajectory from OMN spikes. **a**, Double exponent kernel as fitted from the data, black and gray traces represent the mean and s.e.m over all kernels optimized. **b**, Fast (*τ*_-_ ≈ 5 ms) and slow (*τ*_+_ ≈ 120 ms) time constants of the fitted kernels, shown for all analyzed OMNs. **c**, Actual and predicted eye position during a single trial. Lower panel, zoom in on fixational segment. Note that the predicted eye position is much more variable than the actual eye position. **d**, Distribution across OMNs of correlation coefficients between actual and predicted eye position differences across 350 ms fixational intervals (black). One sided t-test, *p* = 3.8 × 10^−12^ (n=57). Gray, distribution of correlation coefficients obtained from each OMN after shuffling. **e**, For each cell we evaluated what fraction 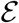 of the variance in actual eye position could be explained by a central noise source. Horizontal axis: 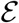. Vertical axis: estimation error of 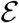 (see also Extended Data Fig.1). Black diamond: weighted average of the results from both monkeys indicates that a large fraction in the variance could be attributed to a central source, 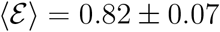 (weighted mean ± weighted error)

## Central source

We next sought to identify whether the weak correlations with eye motion, observed in the noisy activity of single OMNs, are indicative of a dominant central contribution to the motion, upstream of the OMNs. Using the mean of *R* in Fig.2d, 〈*R*〉 = 0.17 ± 0.02 (mean±SEM), we could obtain a lower bound on the average of *R*^2^ across the population of OMNs, by noting that 〈*R*^2^〉 ≥ 〈*R*〉^2^. This bound implies that, on average, at least ~ 2.9% of the variability in eye motion during a 350 ms interval can be predicted based on the activity of a single OMN (SI Notes). This explanatory power of single OMN activity is far too large if the units contribute independently to the eye motion, since there are thousands of OMNs in the primate oculmotor system [23, 24]. We could thus infer that the activity of different OMNs during fixational drift covaries, which provides strong evidence that the eye motion is driven, at least in part, by a common upstream input [25] (see also SI Notes).

The common upstream input to the OMNs could, in principle, be far more dominant in driving fixational drift than implied by the explained variance of single OMNs, because a central contribution to the eye motion would add up coherently downstream of the OMNs, in contrast to the spiking noise of individual OMNs which is independent in different neurons. Therefore, it is not straightforward to assess the upstream input’s contribution to fixational drift based on single neuron recordings. We estimated this contribution by evoking two assumptions: first, that the eye position is linearly driven by the activity of all OMNs – which, in turn, are driven by a common input and additive noise. Second, that the same linear relationship between OMN activity and eye position holds during large eye motions and during fixational drift. Under these assumptions the estimator of eye position, which was fitted to the activity of each OMN during large eye movements, constitutes an unbiased estimator of the common drive to the eye position during fixation. The covariance of the eye translation and the prediction of the estimator can then be used to estimate the common input’s contribution to the variance of eye motion (Methods). Using this approach, we estimated the fraction of the variability in eye motion which is driven by the common input in 350 ms intervals, separately for each OMN. These estimates were noisy, since both the measured and predicted eye trajectories are influenced by sources of noise other than the common input (Methods). The combined results from all OMNs produced a much tighter estimate than obtained from single OMNs, that a fraction 0.82 ± 0.07 (weighted mean± weighted error, Fig.2e and Extended Data Fig.2) of the variance in eye motion during 350 ms intervals is driven by the common input to the OMNs. This key result of our work establishes that most of the eye motion during fixational drift originates in central neural circuits.

## Mean square displacement curves

Even though our analysis above points to a central source for the motion upstream of the OMNs, we sought additional evidence that the correlations between eye motion and OMN activity are not simply a result of the spiking noise in the activity of single OMNs. Such evidence could be obtained by examining how the mean squared displacement (MSD) of the eye motion varies as a function of the time lag. The MSD curves of both monkeys (Fig.3) demonstrated super-diffusive statistics over the entire range of time lags that we examined: on logarithmic axes, the MSD increased steadily with a slope 1 < *α* < 2 (a slope *α* =1 characterizes Brownian motion, whereas a slope *α* = 2 characterizes motion at constant velocity). In addition, the logarithmic slope decreased as a function of the time lag. Both observations are in agreement with measurements in human subjects [9].

**Figure 3:**
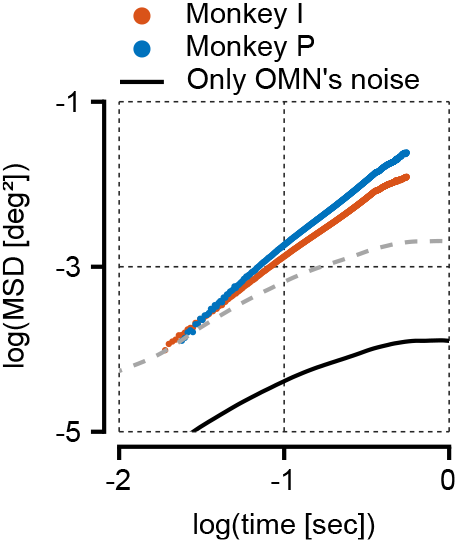
Motoneuron variability can’t explain the mean squared displacement at large time lags. Mean squared displacement (MSD) curves of the recorded trajectories (corrected for the measurement error, blue and red traces), and from simulation (black trace) of 2,000 heterogeneous motoneurons. The black trace flattens towards time lags of order ~1 sec time and can only account for ~ 1% of the total variance at a time lag of ~ 900 ms. Dashed gray trace: translated copy of the black trace along the loagarithmic vertical axis, demonstrates that the slope of the MSD curve generated by filtering of noise due the response of the muscles and mechanics of the eye cannot match the measured MSD curve.

We next estimated the MSD curve of the eye motion that would arise from independent spiking variability of OMNs, driving the occular muscles (Methods). The estimated MSD was too small to account for the experimentally observed variability across all time lags (Fig.3), and at time lags exceeding ~ 0.1 s the predicted MSD was negligible compared to the experimentally observed MSD, in agreement with our previous conclusion that motion is driven upstream of the OMNs.

Furthermore, the predicted logarithmic slope of the MSD curve, generated by the OMN spiking noise, was small at all time lags compared to the measurements, and beyond a few hundred ms the predicted curve saturated, whereas the experimentally measured MSD curve continued to increase steadily (Fig.3). Similar saturation is expected to arise from any form of temporally uncorrelated noise which is fed into the muscle dynamics, at time lags exceeding the characteristic time scale of the muscle response (~ 180ms). Thus, the steep logarithmic slope of the experimental MSD and its non-saturating behavior indicate that the input to the OMNs must itself be characterized by diffusive statistics, with a MSD curve that increases steadily as a function of the time lag, at least up to time lags of order 1 s. This conclusion provides an important insight into the possible underlying mechanism.

## Stochastic diffusion in a memory circuit

OMNs receive their input from the oculomotor integrator, a memory network which is capable of holding the eyes still between saccades by providing steady input to the ocular muscles [20]. Since the horizontal eye position is a continuous variable, the oculomotor integrator maintaining horizontal gaze is commonly modeled as a continuous attractor neural network [26, 27, 28, 29]. It has long been argued on theoretical grounds that in such networks, neural noise can drive diffusive motion along the attractor, which gradually degrades the stored memory [30, 31, 32]. Successes in directly observing this diffusive motion, and especially in relating it to neural mechanisms in specific brain circuits have been scarce and incomplete [33, 34].

The MSD of a diffusive process increases linearly as a function of the time lag. Hence, noise within the ocolomotor integrator has the potential to drive ocular motion with a nonsaturating MSD curve. For this reason, we hypothesized that fixational drift is driven by diffusion within the oculomotor integrator. To put this hypothesis to quantitative test we adapted a model of the goldfish oculomotor integrator [28] to the parameters of the primate visual system (see Methods). Specifically, key parameters that affect the diffusivity [30] such as the number of neurons, their tuning curves, and their spiking variability, were set based on experimental estimates in Macaque (Methods). With these parameters, the output of the network exhibited random diffusion, with a slope of the MSD curve (using logarithmic axes) close to unity (Extended Data Fig.3).

To predict the eye motion statistics, we constructed a mathematical model of central and peripheral contributions to the eye dynamics: the stochastic dynamics of the oculomotor integrator, OMNs, muscles and the ocular plant. We also incorporated in the model a visual feedback mechanism, which can partly correct for the motion of the target due to drift but involves a relatively long delay due to the likely involvement of the cortical or sub-cortical areas [9, 35, 36] (Fig.4a). The output of the integrator was used as an input to a diverse population of OMNs innervating different extra-ocular muscle fibers, with intrinsic spiking noise [37] and with dynamic impact on the muscles which was modeled based on [21] to determine the horizontal eye position.

**Figure 4:**
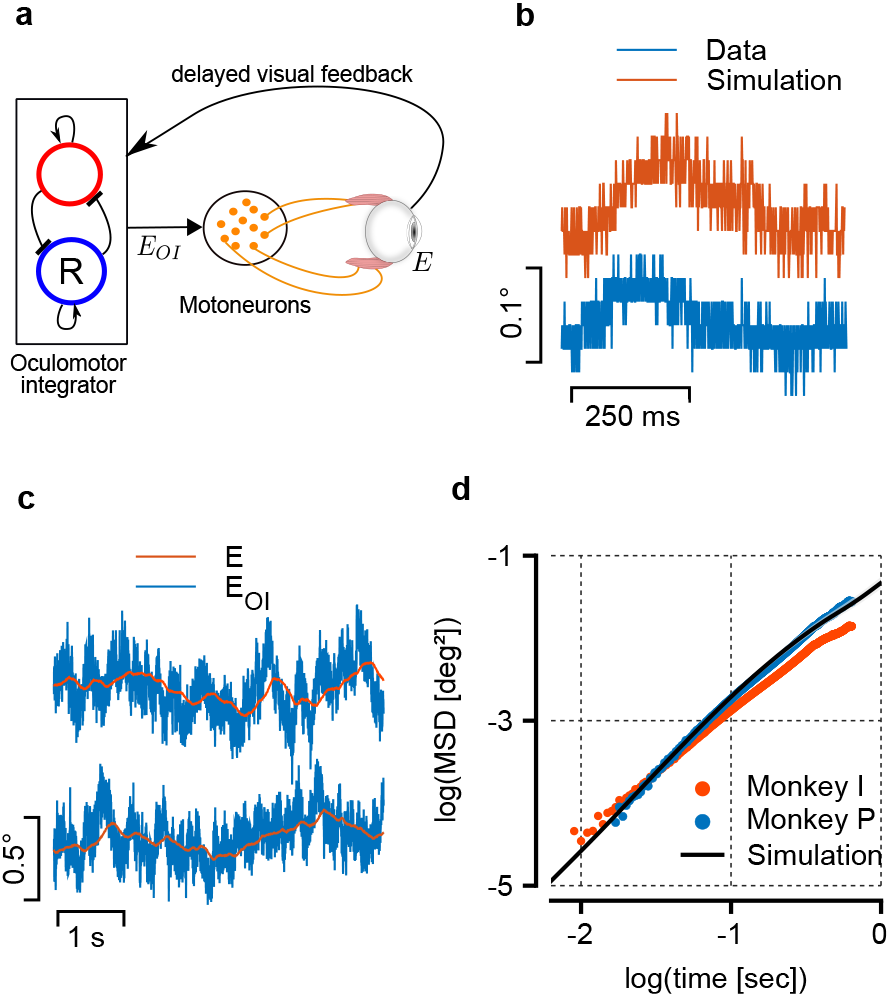
The model. **a**, Schematic illustration of the model: an internal representation of the eye position, *E_OI_* is maintained by the oculomotor integrator and transmitted to the OMNs. OMNs activate the extraocular muscles, resulting in the actual eye position, *E*. The actual eye position serves as a delayed visual feedback to the oculomotor integrator. **b**, Example of simulated and measured eye fixational trajectories. The model generates eye trajectories which are qualitatively similar to the recordings (digitization applied to the simulated trajectory to facilitate the comparison). **c**, Simulated trajectories of *E_OI_*, the internal eye representation (blue), and of *E,* the corresponding actual eye position (red). *d,* Mean squared displacement (MSD) curves, shown using logarithmic axes, as measured from the eye trajectory recordings (blue and red traces) and from simulation of the computational model in a (black).

Numerical simulations of the model using biologically plausible parameters (Methods) generate eye trajectories that resemble measured eye trajectories (Fig.4b). The relationship between the output signal of the oculomotor integrator and the actual eye position is demonstrated in Fig.4c: the oculomotor integrator output is noisy due the spiking variability of oculomotor integrator neurons, whereas the actual eye trajectory is smoother, due to the mechanics of the muscles and the eye ball. When examined at a coarse scale of hundreds of milliseconds, the eye position follows the output of the oculomotor integrator. Importantly, the simulated MSD curve of the eye position (black trace, Fig.4d) increases steadily as a function of the time lag at time scales exceeding ~ 100ms due to the diffusive dynamics of the oculomotor integrator output.

The main consequence of the visual feedback mechanism in the model is to decrease the logarithmic slope of the MSD curve at large time scales, which is otherwise a bit too steep compared to the experimental measurements (Extended Data Fig.5, see also SI Notes). On the other hand, the visual feedback mechanism has little influence on the slope of the logarithmic MSD curve at shorter time scales (Extended Data Fig.4a), and is unlikely to affect this feature due to the large synaptic delays involved. Therefore, we propose that the reason for the super-diffusive statistics of fixational eye drifts lies in the diffusive dynamics within the oculomotor integrator, and is largely independent of visual feedback mechanisms.

In summary, with a few parameters – most of which were chosen based on known features of the primate oculomotor system (Methods), the model produces simulated MSD curves that match experimental observations very well, both in magnitude and in shape (Fig.4d and Extended Data Fig.6).

## Discussion

We showed for the first time that fixational drift is correlated with neural activity, and identified the main source of the motion in central neural circuitry upstream of the OMNs. The statistics of the motion provided an important clue on the identity of the upstream drive, pointing to noise-driven diffusion in the oculomotor integrator as a likely source of the motion. Theoretical modeling, constrained by the physiology of the primate oculomotor system, provided further support to this hypothesis since it accounted for the magnitude and detailed statistics of the motion. Taking the view that the oculomotor integrator is a short-term memory network [26, 28, 29], we thus propose that fixational drift offers direct observation of diffusive dynamics in continuous attractor networks, and an arena for probing mechanistically how noise affects the storage of continuous parameter memory in neural circuits.

Both intrinsic variability and noisy inputs can drive stochastic diffusion in continuous attractor networks. We assumed that the noise is intrinsic to neurons within the oculomotor integrator network, and this assumption accounted well for the amplitude of the motion. However, we only have approximate estimates for some parameters, such as the number of neurons in the network. Hence, we cannot rule out that the diffusive dynamics in the state of the oculomotor integrator is driven in part by noisy premotor inputs to this network, such as those arising from vestibular, optokinetic, and vergence signals.

Our focus on noise in the occulomotor integrator as a drive of diffusive input into the OMNs does not imply that fixational drift is unaffected by additional mechanisms, acting within the visual-motor pathway. The incorporation of a visual feedback mechanism in our model demonstrates, indeed, that a visual feedback loop can modulate the statistics of the motion at time lags exceeding *~* 100 ms – but is unlikely to influence the slope of the MSD curve at shorter time scales, due to the large synaptic processing delays involved. We note also that the strength and dynamics of modulation by visual feedback are likely affected by the salience and structure of the visual stimulus, in accordance with the observations that the detailed statistics, but not the superdiffusive nature, of fixational drift is influenced by the visual task [5, 12, 36, 38, 39, 40, 41].

Finally, fixational drift is highly consequential for visual perception in the fovea, even though its amplitude is tiny compared to saccades [5, 10, 11, 12, 13, 15, 16]. Detailed understanding of the mechanisms underlying fixational drift is likely to advance the research on its functional consequences, by opening up the possibility to examine whether specific parameters of the oculomotor pathway, such as the variability of neural activity within the oculomotor integrator and the dynamics of the muscle response, are tuned to optimize visual function.

## Methods

### Behavioral task and recordings

We have reanalyzed data reported in published studies [2, 3, 42]. Data were collected from two male rhesus macaque monkeys (Macaca mulatta) that had been prepared for behavior, and neural recordings were obtained using techniques described in detail previously. All procedures had been approved in advance by the Institutional Animal Care and Use Committee at UCSF, where the experiments were performed. Procedures were in strict compliance with the National Institutes of Health Guide for the Care and Use of Laboratory Animals. Briefly, glass-coated platinum-iridium electrodes were lowered into the brainstem to record from neurons. The abducens nucleus was distinguished by the characteristic singing activity associated with ipsiversive eye movements. We identified OMNs, as opposed to internuclear neurons (INNs) based on the criteria used in [43], Extended Data Fig.2. Visual stimuli appeared on a monitor at a distance of 30 cm from the monkey’s eye. Targets were bright 0.6° circles on a dark background. We recorded neural activity during pursuit of step-ramp target motions [44]. At the start of each trial, the monkeys had a second to acquire fixation on a stationary target. They were then required to fixate for an additional 500-700 ms within a 2°-3° square window. The target then displaced to a location eccentric to the position of gaze (step), and immediately began moving toward the fixation point (ramp).

### OMN firing rate estimators

We obtained estimators 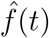 of the OMN firing rates based on the eye trajectory as follows. The estimators were expressed as a linear combination of the eye position, velocity, and acceleration [20, 45, 46, 47]:

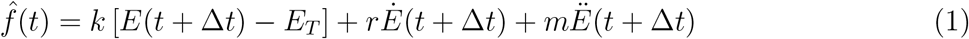

where the parameters *k, r, m*, Δ*t*, and *E_T_* were chosen to fit simultaneous measurements of the OMN firing rate during the full extent of each trial, including the fixational and smooth pursuit components. The firing rate *f*(*t*) was extracted from the spike train as the inverse of the inter spike interval [20], and was thus taken to be constant between successive spikes. We discarded trials in which the measured firing rate went below 20 Hz [21], since equation(1) only holds above the OMN threshold. The parameters were chosen to minimize the loss function

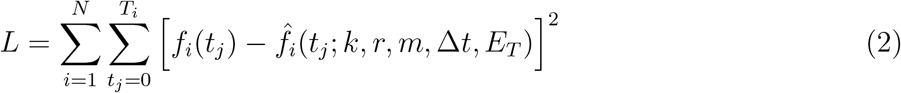

where *T_i_* is the duration of the *i*-th trial, *t_j_* ∈ [0,*T_i_*] are discrete time samples within a trial, and *N* is the total number of trials. To generate an estimate to the local eye acceleration we smoothed the velocity signal using a Savitzky-Golay filter of order 3 and 21 ms length [48] and performed a two-sided numerical differentiation. To generate a firing rate prediction for each of the hundreds of trials per cell we followed a “leave one out” scheme: we fitted the cell parameters to all but one trial, and used these parameters to generate the firing rate prediction for the left out trial.

### Extracting the fixational segments

Microsaccade onsets and endings were detected using the eye motion in 2d (horizontal and vertical components), by applying a combination of thresholds for the magnitude of the velocity (10 deg/sec) and the magnitude of the acceleration (1000 deg/sec^2^). The fixation segments used for further analysis started 30 ms after termination and ended 30 ms before initiation of the microsaccades. Finally we verified manually that the fixational segments didn’t contain any microsaccades.

### Firing rate correlation coefficients

Correlation coefficients in the firing rate representation (Fig.1) were calculated by generating two vectors of the actual and predicted firing rates from all trials available per cell. Each vector was generated as follows: for each fixational segment we calculated the firing rate, either actual or predicted, and substracted from it the mean firing rate during that fixation. Finally we concatenated contributions from all fixational segments and calculated the Pearson correlation between the predicted and actual firing rate vectors.

### Model inversion

Eye position was predicted from the spiking activity by causal filtering:

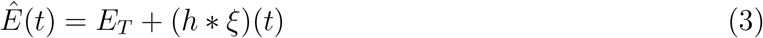

Here, *ξ*(*t*) is the spike train emitted by the cell:

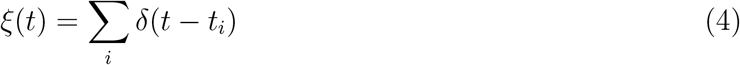

where *t_i_* is the timing of the *i*-th spike and *h*(*t*) is the kernel of the linear filter that represents the inverse of the relationship in equation(1):

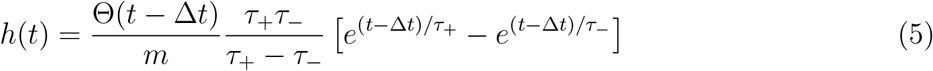

where

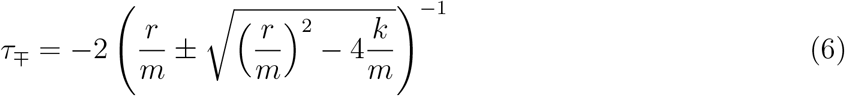

and Θ is the Heaviside step function.

### Eye position correlation coefficients

Correlation coefficients between the measured eye position and its prediction based on single OMN spike trains (Fig.2d) were evaluated as follows. First, equation(3) was used to generate a prediction *Ê*(*t*) of eye position. We discarded fixational segments shorter than 350 ms and broke longer segments into non overlapping 350 ms fixational segments. For each fixational segment we calculated the difference in the measured eye position, *δE* = *E*(*t* = 350 ms) – *E*(*t* = 0 ms), across the fixational segment, and the corresponding prediction *δÊ* = *Ê*(*t* = 350 ms) – *Ê*(*t* = 0 ms). Finally, we calculated the Pearson correlation coefficient between all predicted and measured eye position differences

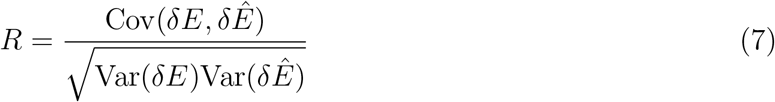

In addition we calculated the Spearman rank correlation between these two sets of eye differences which also demonstrated significant correlation (p=2.5 × 10^−12^, Extended Data Fig.7).

### Standard error of the estimates for the covariance and *R*

The standard error of the covariance estimate (used to calculate the errors in Fig.2e, Extended Data Fig.1) is given [49] by

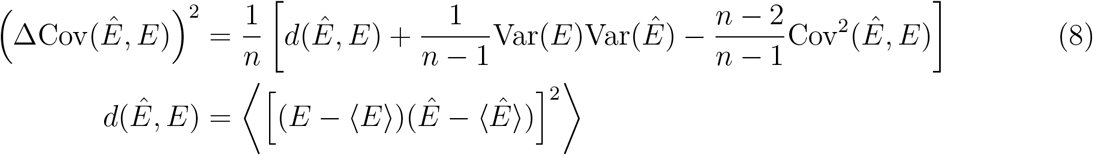

where *n* is the number of fixational segments used to evaluate the covariance.

The standard error of the estimate for *R*, equation(7) was evaluated as follows:

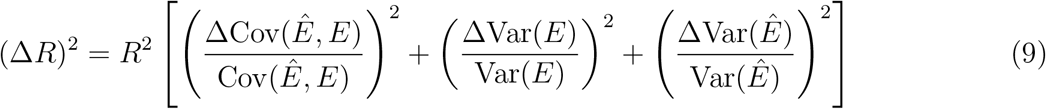

where ΔVar(*E*) and ΔVar(*Ê*) are the standard errors of the estimates for Var(*E*) and Var(*Ê*), obtained using expressions similar to equation(8).

### Estimated contribution of central source to measured eye position

To estimate the contribution of a central source to the measured eye position, we evoke the following assumptions. We assume that the measured eye position *E* can be written as:

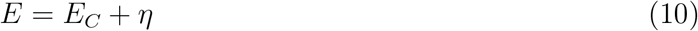

where *E_C_* is a contribution arising from sources upstream of the OMNs, and *η* represents contributions arising from the OMNs as well as other sources of noise downstream of the OMNs. We assume that *η* and *E_C_* are uncorrelated.

Similarly, we assume that the estimate of eye position *Ê*, obtained from the spike train of a single OMN can be written as

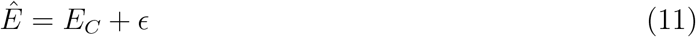

where *ϵ* represents the error of the estimator and is assumed to be uncorrelated with *E_C_*. Thus, we assume that the estimator, which was fitted to measurements obtained during large eye motions, remains unbiased during fixational intervals. Note that during fixation the variance of *ϵ* is large compared to that of *E_C_*, which is the main reason why the correlation coefficient between *E* and *E_C_* is small compared to unity.

Our goal is to estimate what fraction of the variance in the measured eye position, *E,* is due to the central source, *E_C_*, over a fixational segment (in our analysis we took segments of duration 350 ms):

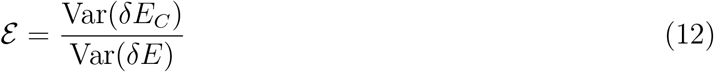

where *δ_x_* represent the difference between the value of a variable *x* at the end of the interval and its value at the start of the interval. Under the assumption that the noise terms η and e are uncorrelated,

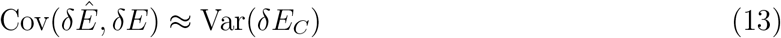

In fact, there is a very weak correlation between *η* and *ϵ* which arises from the fact that *η* includes the noise contributions from all the OMNs, and e represents the contribution of noise to the estimate error from a single OMN. This correlation however, is of order 1/*N_m_* where *N_m_* is the number of OMNs, and we can safely neglect it, as corroborated also from our separate estimate of the OMN contribution to fixational drift (Fig.3). Therefore, our estimate of 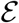 is given by

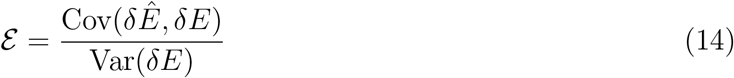

The standard error in our estimate of this quantity is dominated by our ability to measure the covariance from a limited number of trials. The estimated errors in Fig.2d were thus calculated as

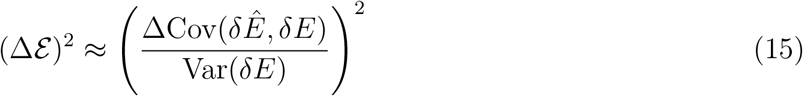

Note, that ideally in equation(15) we would like to include in the denominator the variance of the true eye position and not the measured eye position. However over intervals of duration 350 ms the measurement error of the change in the eye position is negligible compared to the actual motion, and is inconsequential for our estimate of the relative contribution of the central source to the motion.

Assuming that the errors in our estimates of 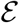 are normally distributed, we calculated a weighted mean of this quantity across all cells from both monkeys as follows

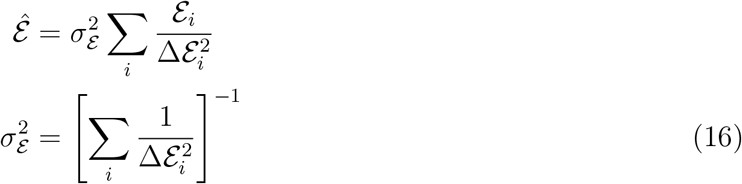

where 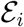 and 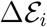 are the estimated value and the estimated standard error of 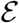 from the *i*’th cell. We used a weighted mean, and not a simple average over all cells, because of the large variation in 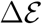 across cells, which was due to differences in various factors such as the number of trials and the structure of the tuning curves.

### Empirical MSD curves

The measurements of the eye position from the search coil are corrupted by a measurement error of variance ~ 10^−3^ deg^2^. This is evident when plotting the MSD curve of the eye movements, where the measurement error gives rise to a flat MSD curve (when using logarithmic axes) up to time scales in which the diffusion becomes dominant. A similar estimate for the measurement noise was obtained by recording from a coil fixed in space. Therefore, in Figs. 3 and 4d we subtracted the variance of the measurement noise to recover the MSD statistics of the eyes themselves. The MSD curve prior to noise subtraction is shown in Extended Data Fig.8.

### Model of fixational drift

Our model of fixational drift (Fig.4a) consists of three stages. First, an internal signal of the desired eye position is generated and held in the memory network of the oculomotor integrator. In the second stage this signal is conveyed to a population of spiking OMNs. The synaptic output of each OMN is translated into an actual eye position using equation(1). Finally, The actual eye position is taken to be an average over all OMNs.

#### Oculomotor integrator network

The oculomotor integrator network model is based on the model proposed in [28] for the neural network that determines the horizontal eye position in goldfish, adapted to the parameters of the primate oculomotor system. Briefly, the network consists of two populations, of which one is more active when the eye is directed to the left and the other is more active when the eye is directed to the right. Each population forms excitatory synaptic connections to itself and inhibitory synaptic connections to the other population. The connectivity is tuned such that there is a continuum of steady states, representing a continuum of stable eye locations, and such that the firing rates of the single neurons fit the experimentally observed tuning curves across the full range of eye positions.

We adapted the model of [28] in the following ways. First, we introduced noise by using spiking neurons, with a CV of the inter spike intervals of ~ 0.22 [3, 50] as described below. The oculomotor integrator network dynamics are thus described in our model by the following equations

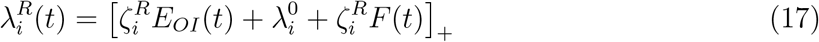

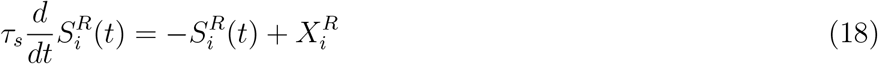

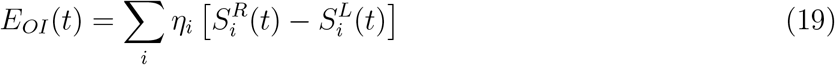

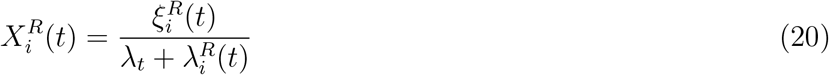

In equation (17) 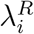 is the firing rate of neuron *i* from the right population, and [*x*]_+_ = max (*x*, 0). In equations(18-19) 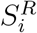 is the synaptic output generated by neuron *i* of the right population. We similarly denote by 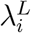 and 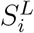 the firing rate and the synaptic output of neuron *i* from the left population (see below). The spike train of neuron *i* from the right population is denoted in equation(20) by 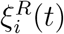, and *τ_s_* in equation(18) represents the characteristic time scale of post-synaptic currents. As in [28], the influence of spikes on the synaptic current is nonlinear, through the relationship between and 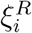, and 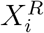 equation(20), where *λ_t_* is a characteristic firing rate, in our case *λ_t_* = 80 Hz. Note that if 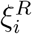 is replaced in equation(20) by 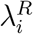, the dynamical equations become identical to those of [28]. In equation(17) *F*(*t*) represents a visual feedback signal, which is determined as described below.

The variable *E_OI_*(*t*) appearing in equation(17) represents an internal readout of the eye position from the neural activity, which is a linear function of the synaptic activities with weights *η_i_*. This quantity also serves as the eye position readout from the network. The slope of the tuning curve of neuron i from the right population is denoted by 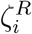 and is positive, and 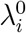 is an offset. In those neurons that are above threshold at central gaze (i.e. eye position zero), 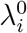 is equal to the firing rate at central gaze.

For the left population, equations analogous to (17), (18), and (20), are obtained by replacing the superscripts *R* and *L.* We assumed that for each neuron in the right population there is a matching neuron in the left population whose tuning curve is identical up to reversal of sign of the eye position. Hence, the slopes of the tuning curves in the left population are set as 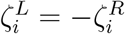.

The tuning curves were enriched from [51] by randomly choosing slopes from the values *ζ_i_* measured in the goldfish, and modifying them by sampling new values from a uniform distribution in the range of 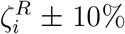. In addition, we adapted the slopes by multiplying them by a fixed prefactor (identical for all oculomotor integrator neurons), which was chosen to match the typical firing rates in primates during straight ahead gaze, ~ 80 Hz [52, 53, 54]. The weights *η_i_* were determined by an optimization process as in [28], ensuring that the system has an approximate continuum of fixed points in which the eye position variable *E_OI_*(*t*) spans the range of −20 to 20 degrees. The precise values of 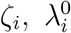, and *η_i_* are listed in SI Data.1.

#### OMN activity and actual eye position

The internal eye representation generates activity of the OMNs according to the following dynamics

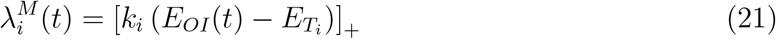

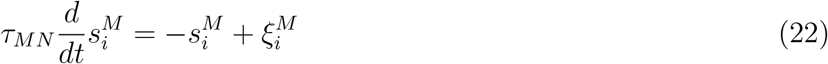

where 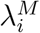, the firing rate of OMN *i*, is assumed to be determined linearly and instantaneously from the synaptic readout *E_OI_*(*t*) of the oculomotor integrator network. We incorporate noise in activity of the OMNs by generating a stochastic spike train 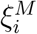 with rate rate 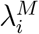 and with a CV of ~ 0.07 as described below. Each OMN generates a synaptic output 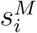 which is determined by equation(22).

Each OMN is assumed to innervate an extraocular muscle fiber and to contribute a signal *E_i_* to the actual eye position, which can be expressed as a convolution of *E_OI_* with a double-exponential kernel or, equivalently, as the solution of the differential equation

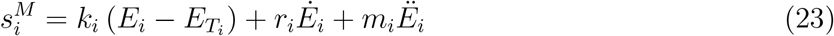

with parameters *k_i_, r_i_,* and *m_i_*. Note that equations (21-23) are set up such that when the eyes are still *E_i_* = *E_OI_* up to stochastic fluctuations (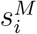 is equal at steady state to 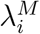, up to stochastic fluctuations, due to equation(22)).

Finally, the position of the eye is taken to be an average over all the OMN contributions:

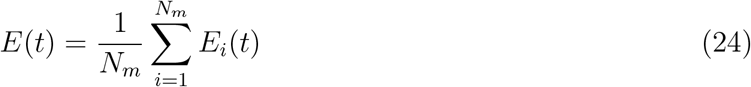

When the eyes are still and at steady state the actual eye position *E* matches the internal representation of the oculmotor integrator *E_OI_* [55, 56].

The parameters *k_i_, r_i_, m_i_*, and *E_T_i__* of the OMNs were set as follows. First, the eye position threshold *E_T_* was sampled uniformly in the range (–45°)–(–5°), with mean eye threshold of –25°, similar to [20], we discarded neurons with eye thresholds above –5° since their contribution to eye position during straight ahead gaze is negligible. Second, we assumed an approximate linear relationship between *k, r* and *E_T_*, as observed in [21]. Thus, *k_i_* and *r*_i_ were set as

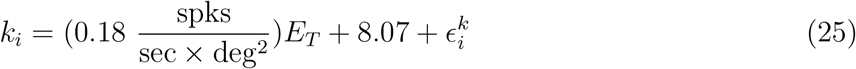

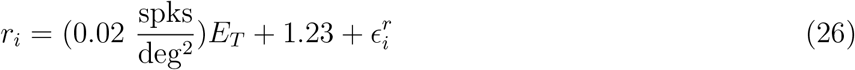

where the coefficients of the linear relationships were set based on [21], and where 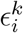 and 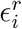 are zero-mean, normally distributed random variables whose variance was chosen to obtain the same dispertion as experimentally observed in [21] (correlation coefficients *R* = 0.81 for *k* and *R* = 0.67 for *r*). In addition we imposed hard constraints *k* ≥ 1.1, *r* ≥ 0.25 according to [20]. The typical values for *k*, and *r* obtained from this procedure matched the values reported in [20]. For the acceleration coefficient, *m*, we randomly sampled values according to typical values reported in [20] while verifying that *m* < *r*^2^/(4*k*). The precise parameters values of *r, k*, and *m* are given in SI Data.2 and illustrated in Extended Data Fig.9.

Our simulation included *N* = 20, 000 neurons in the oculomotor integrator, *N_m_* = 2, 000 OMNs, synaptic time constants of *τ_s_* = 10 ms, and visual feedback amplitude *A* = 0.035 (see below). We used a set of parameters that is biologically plausible, but note that other combinations of parameters can produce similar results. Specifically, several key parameters determine together the amplitude of the MSD curve: the number of neurons in the oculomotor integrator, their synaptic time constant, firing rate, non-linearity, and variability [30].

#### Visual feedback

We assume that the visual feedback during fixational drift is based on motion of the target on the retina, rather than on attempt to fix the absolute position of the target [40]. Therefore, the visual feedback signal *F*(*t*) in equation(17) is proportional to an estimate of the target angular velocity relative to the eye direction in the recent history. The estimate of velocity is generated using a signal that arrives to the oculomotor integrator with a delay *τ_d_* ≈ 70 ms [9]:

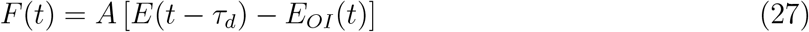

where *E*(*t*) is the actual gaze direction established as described above.

#### Sub-Poisson spike trains

Both oculomotor neurons and motoneurons are sub-Poisson with CV of the inter-spike interval < 1. In order to take this into account in our model we used spike thinning by first generating Poisson spikes at a rate equal to the desired firing rate, multiplied by a factor *M.* From the resultant spike train we then used every *M*’th spike. This procedure keeps the average firing unchanged, but reduces the CV by a factor 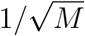. When the firing rate is constant, this procedure is equivalent to sampling inter-spike intervals from a Gamma distribution. For oculomotor neurons we assumed CV~ 0.22, for the OMNs we assumed the CV is linear with the mean inter-spike interval, (ISI) as observed in [37]

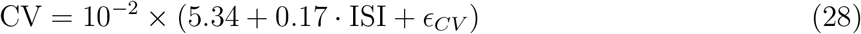

where the ISI, is measured in ms, and *ϵ_CV_* is a zero-mean, normally distributed random variable whose variance was chosen to obtain the same dispersion as experimentally observed in [37]. Note that these values are based on the empirical distribution of inter-spike intervals during fixational intervals and could therefore be influenced, in principle, by the small motion of the eye during fixational drift. However, we show in SI Notes (Contribution of oculomotor state to the CV of OMNs) that this influence is negligible.

## Extended data figures and tables

**Extended Data Fig. 1:**
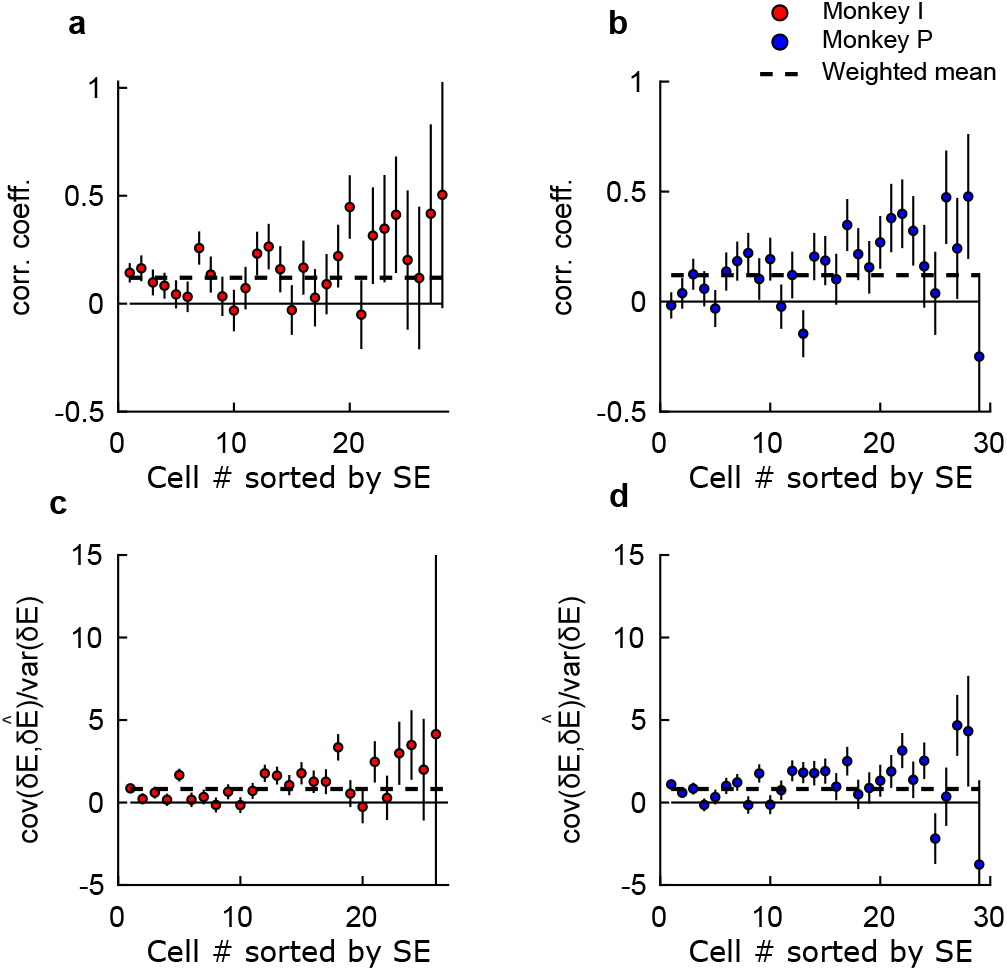
Correlation coefficient and 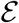 sorted by their SE. **a**, Correlation coefficients as measured for each OMN in monkey I. **b**, Same as in (a) but for monkey P. **c**, The estimated fraction of variance in the measured eye position which is due to the central source (see Methods) measured in monkey I. For clarity two cells with 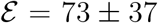, 44 ± 41 are not shown. **d**, Same as in (c) but for monkey I. In all panels, cells are ordered by the standard error of the estimates.

**Extended Data Fig. 2:**
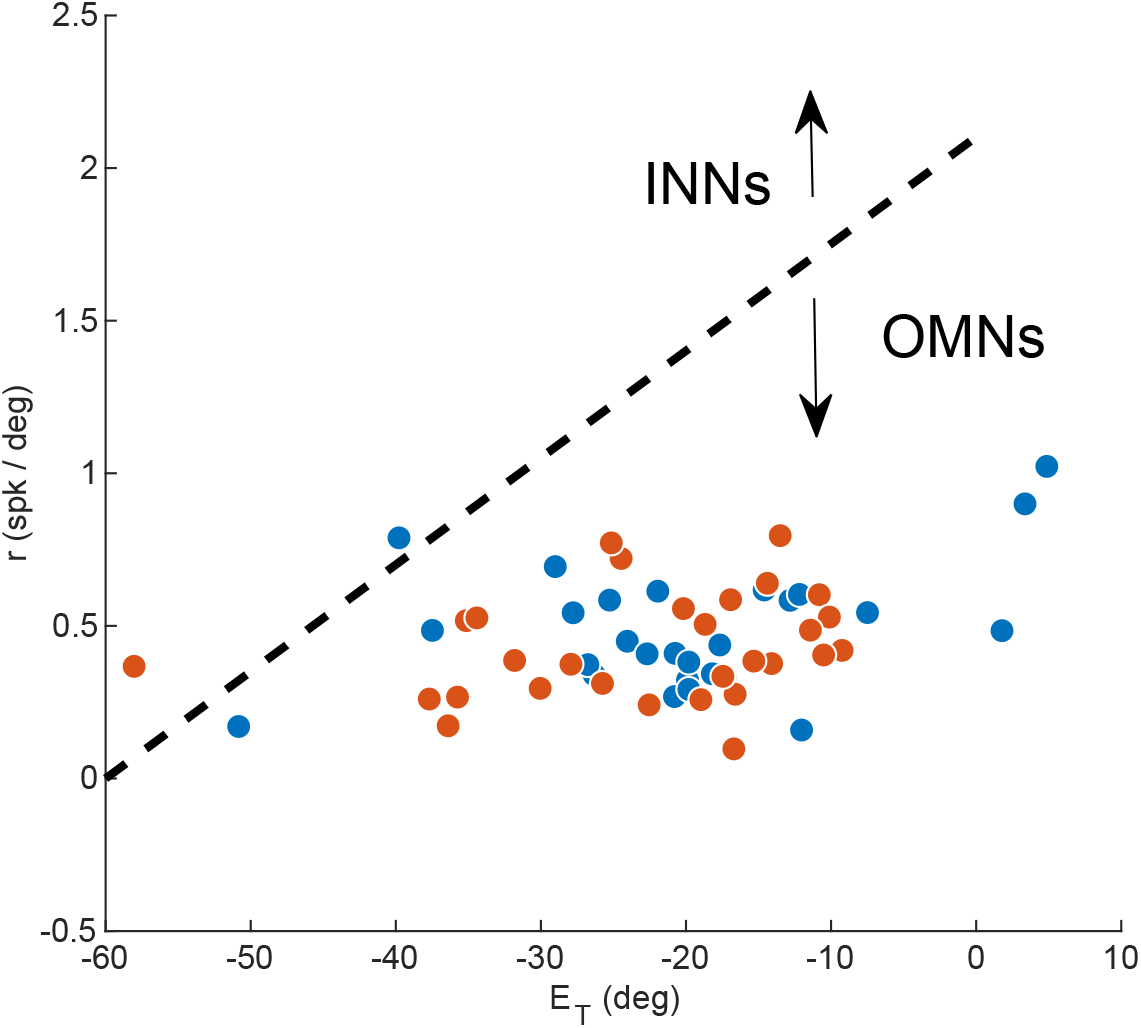
Identifying OMNs vs. INNs. Recorded cells in our data set were identified as OMNs vs. internuclear neurons (INNs) using a criterion based on tuning curve parameters, as in [43] (dashed black trace): the eye position threshold *E_T_* and the eye velocity sensitivity *r*. (Two cells which were close to the separating criteria defined in [43] from above were included in the analysis. The inclusion of these cells did not qualitatively affect any of the results.)

**Extended Data Fig. 3:**
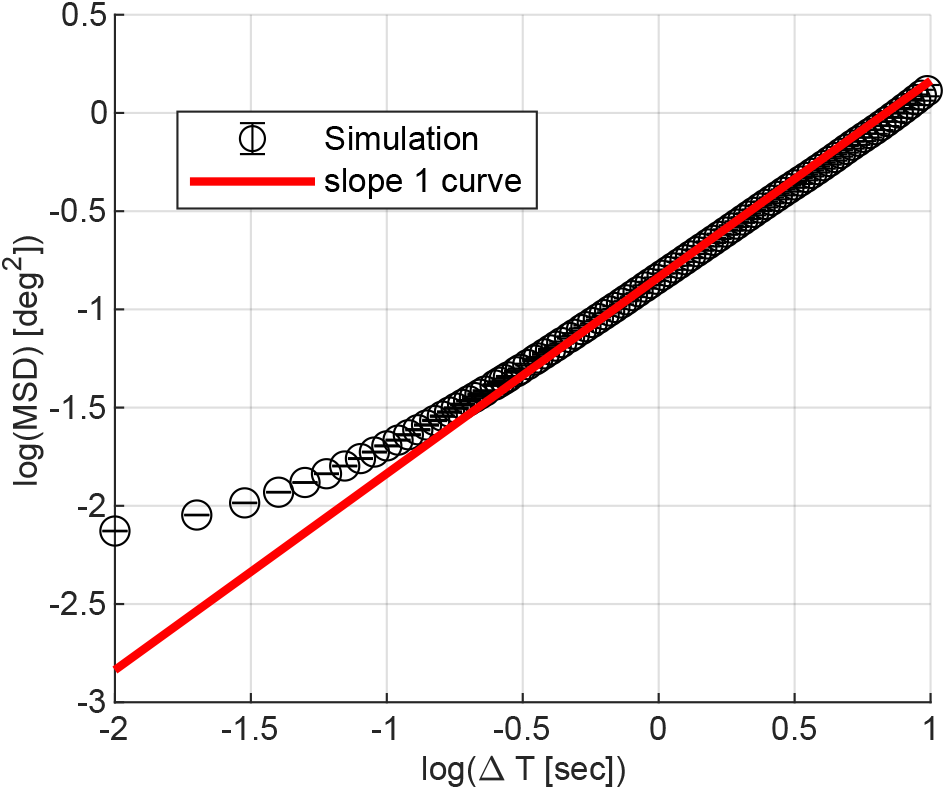
MSD of the simulated oculomotor integrator output. MSD curve of the model oculomotor integrator output, measured in simulations (visual feedback is not included in these simulations). The logarithmic slope of the MSD curve approaches unity at time lags larger than the synaptic time constant, ~10 ms, as expected for a noisy continuous attractor network [30].

**Extended Data Fig. 4:**
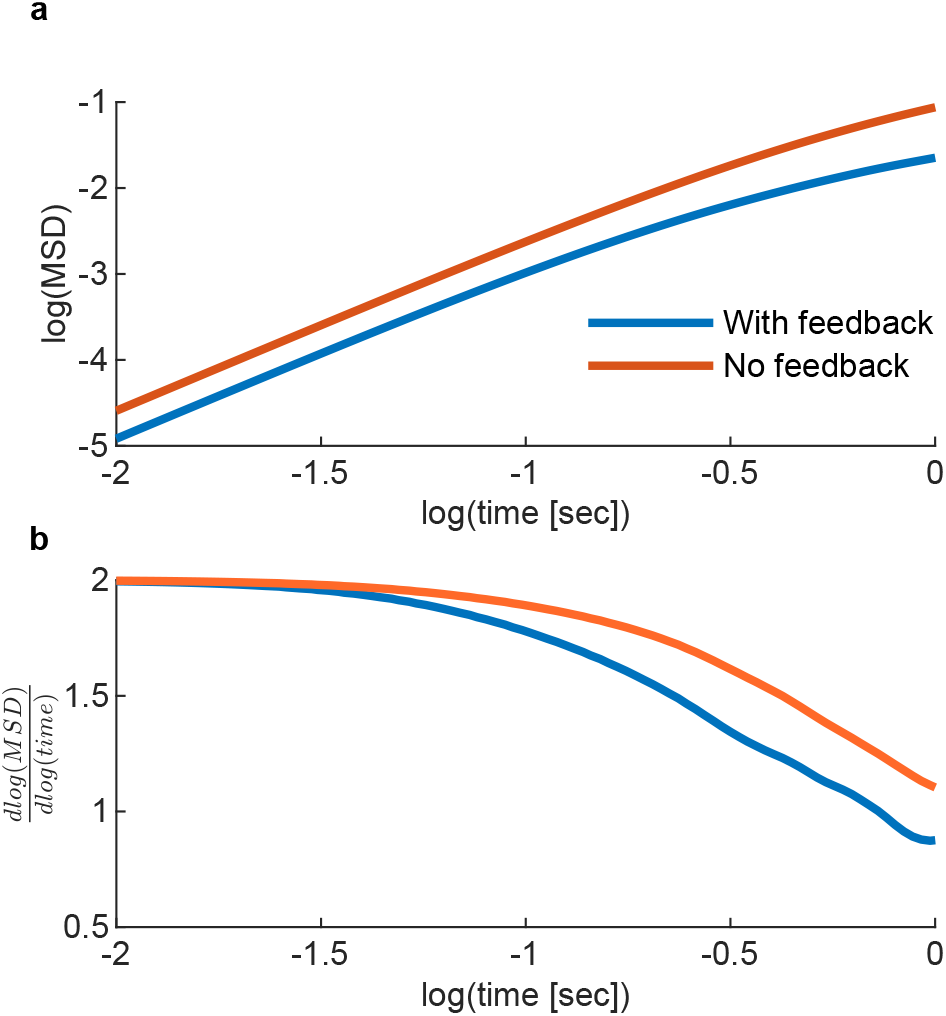
Visual feedback effect on the MSD curve. The effect of visual feedback in the theoretical model (shown without spiking noise). **a**. Feedback decreases the amplitude of the MSD at all time scales, but more significantly at Δ*t* ≳ *τ_f_*. **b**. Feedback decreases the logarithmic slope of the MSD curve at Δ*t* ≳ 100 ms

**Extended Data Fig. 5:**
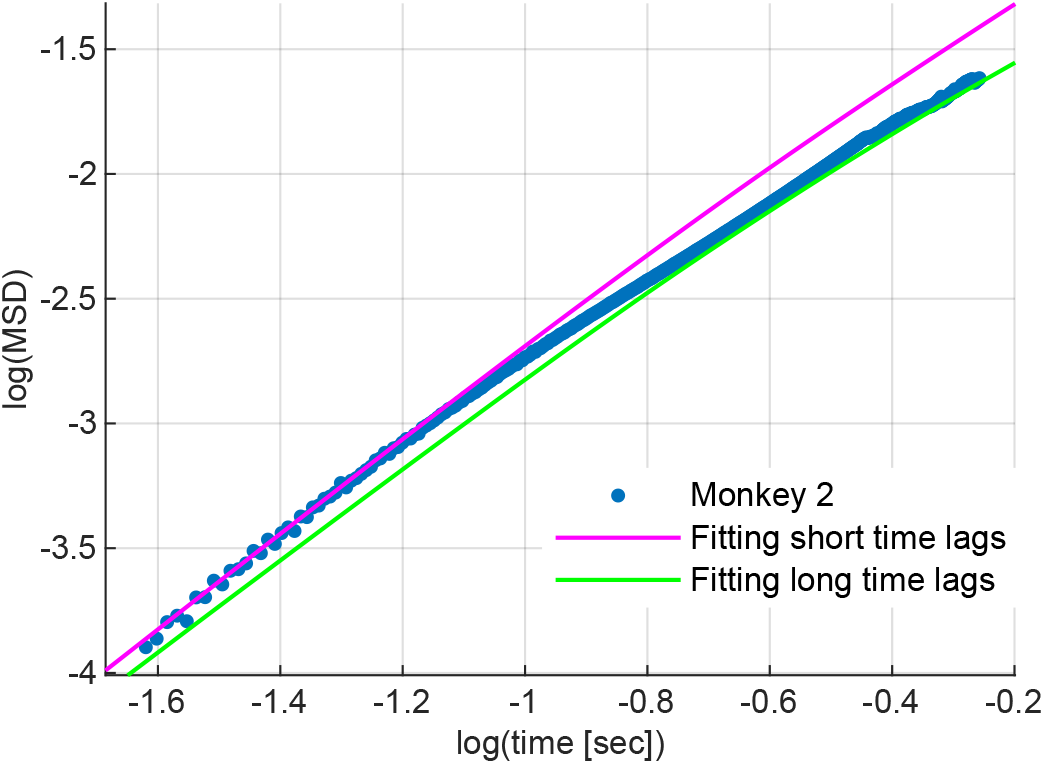
Fitting the data in a model without visual feedback. Without the visual feedback mechanism, the model cannot fit the measured data precisely: it either overestimates the MSD at large time lags or underestimates the MSD at short time lags. Fitting to the behavior at short time scales (magenta) was obtained with *N* = 20,000 oculomotor integrator neurons with CV~ 0.16, and *N_m_* = 1, 800 OMNs. Fitting the behavior at long time scales (green) was obtained with *N* = 30, 000 oculomotor integrator neurons with CV~ 0.16, and *N_m_* = 1,000 OMNs.

**Extended Data Fig. 6:**
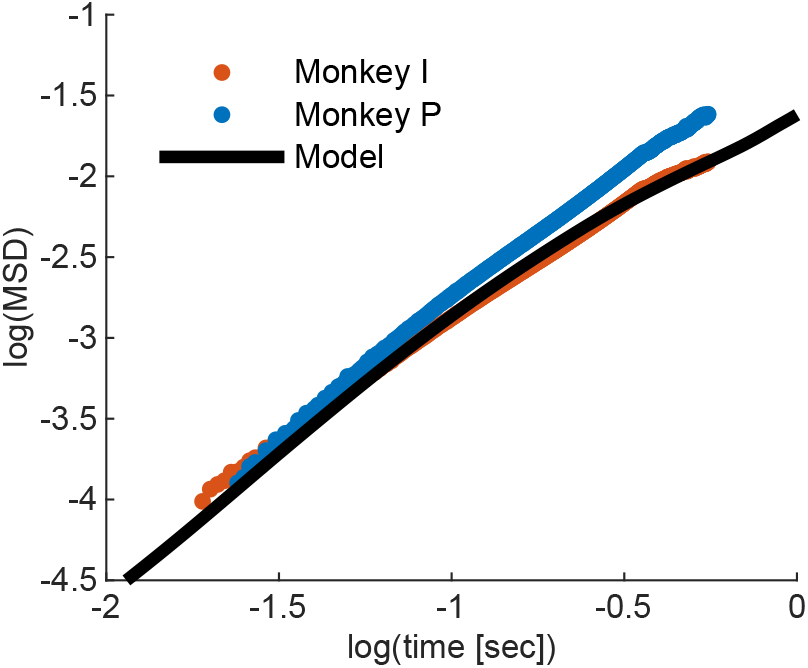
Fitting the MSD curve of monkey I. Here we demonstrate that a precise fit for the MSD curve of monkey *I* can be obtained with relatively minor modifications to the parameters from the values used in Fig.4d: (1) The number of OMNs was decreased from 2,000 to 1,000. (2) The OMN CV was determined from the linear relation 7.0 + 0.18 × *ISI*. (3) The CV of oculomotor integrator neurons was changed from ~ 0.22 to ~ 0.17. (4) Feedback strength parameter *A* in equation(27) was increased from 0.035 to 0.05.

**Extended Data Fig. 7:**
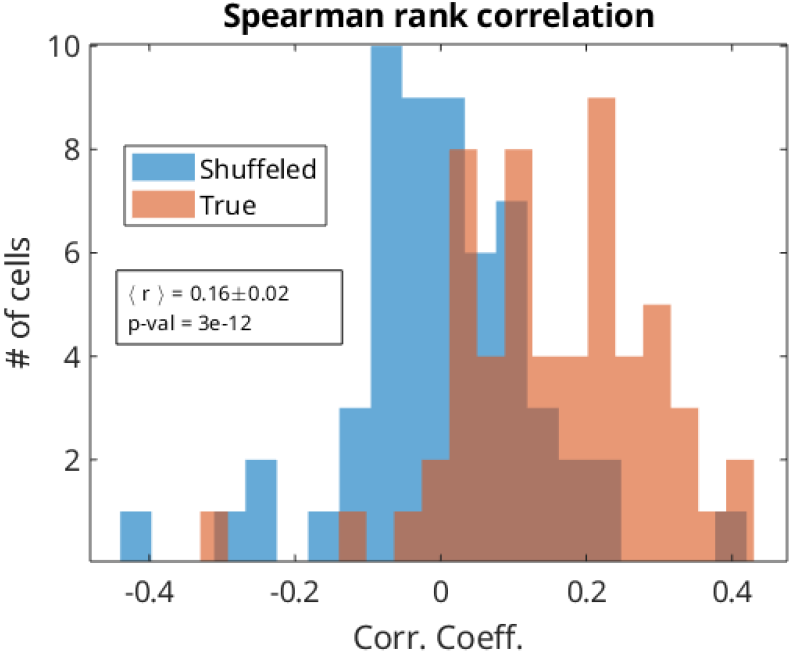
Distribution of Spearman correlation coefficients. The nonparametric correlation coefficient between *δE* and *δÊ* demonstrates a positive and significant mean value of 0.16 with one sided t-test p-value 3 × 10^−12^. (compare with Fig.2d)

**Extended Data Fig. 8:**
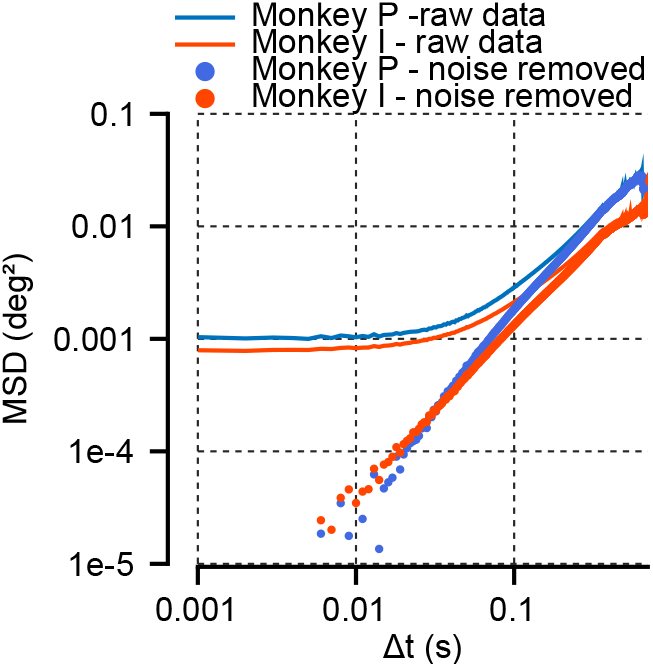
Removal of measurement noise from the MSD curve. The measured MSD before and after measurement noise removal. Saturation of the MSD at short time lags towards ~ 10^−3^ deg^2^ is indicative of the measurement noise variance.

**Extended Data Fig. 9:**
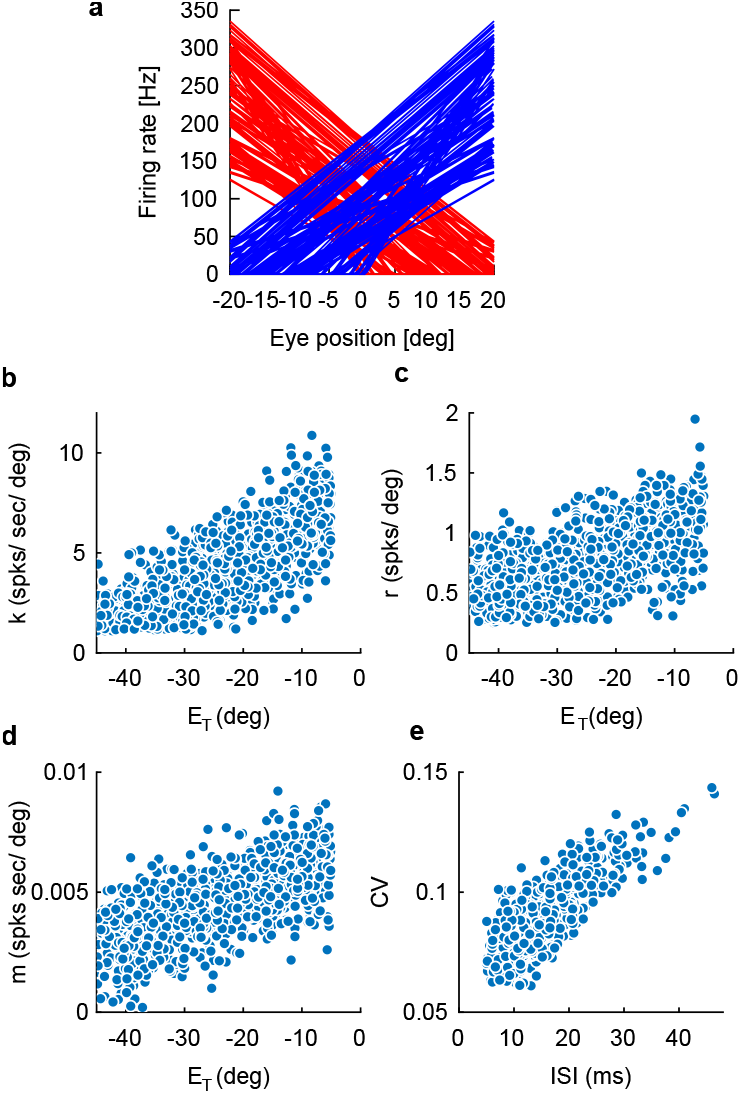
Parameters of the computational model. Visualization of the parameters used in the computational model. **a**, Tuning curves of neurons in the OI, red (blue) color corresponds to left (right) populations. For clarity only 100 representative tuning curves are shown from each population. **b,c,d**, The parameters which define the OMNs response as defined in equation(1). **e**, The CV of each OMN at straight ahead gaze.

**Extended Data Fig. 10:**
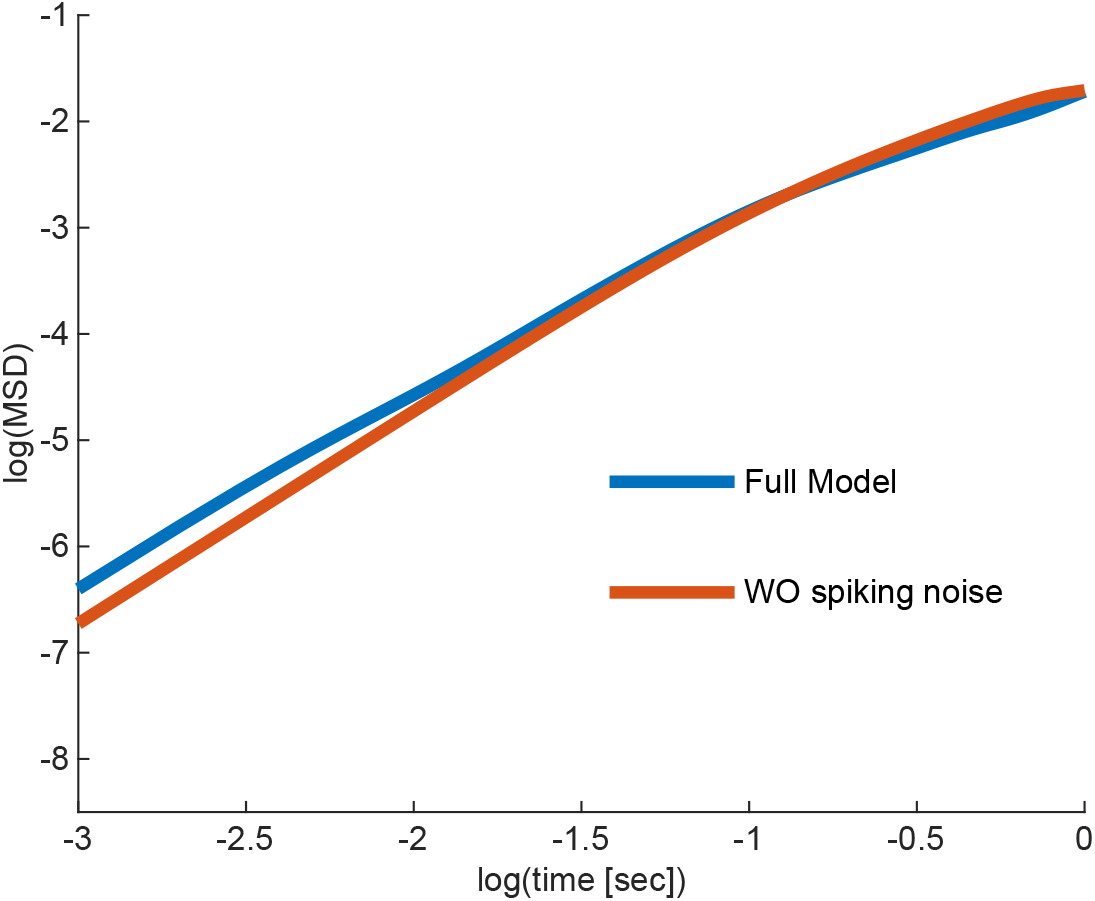
Spiking noise effect in the computational model. MSD curves generated by the full model, and by a model which excludes spiking noise, both in the output of the oculomotor integrator and in the OMNs (stochastic spiking was still included in the internal dynamics of the oculomotor integrator to drive diffusion). The spiking noise effect is negligible at time lags > 50ms, and contributes significantly to the MSD at smaller time lags, up to about half of the variance at a time lag of 1ms.

## Supplementary information

**SI Notes** The explanatory power of a single OMN. Contribution of oculomotor state to the CV of OMNs. Contributions of peripheral noise and feedback to the eye motion.

**SI Data.1** Parameters of the simulated oculomotor integrator network. The first column corresponds to the slopes of the neuronal tuning curves, ***ζ***, the second to the offset at straight head gaze, **λ**^0^ and the third to the optimized values, ***η**.*

**SI Data.2** Parameters of the OMNs in simulations. The first column corresponds to the slopes of the neuronal tuning curves, *k,* the second to the eye velocity sensitivity, *r,* the third to the eye acceleration sensitivity, m, the fourth to the tuning curve threshold, *E_T_*, and the fifth to the CV of each OMN.

## Acknowledgements

This research was supported by the Israel Science Foundation grant No. 1733/13 and grant No. 1745/18, and in part by the Israel Science Foundation grant No. 1978/13. We acknowledge additional support from the Swartz foundation and from the Center for Brains, Minds and Machines (NS), the Israel Science Foundation grant No. 380/17 and the European Research Council grant No. 755745 (MJ), and the Gatsby Charitable Foundation (YB).

## Supplementary Notes

### The explanatory power of a single OMN

In the main text we concluded that OMNs share a common input, since each OMN can explain a relatively large fraction of the variance in the measured eye position. Here we provide further details on the reasoning that leads to this conclusion.

First, consider a predictor of the eye motion, which is based on the activity of a single OMN. The coefficient of determination *R*^2^ between the predictor and the actual motion is equal to the fraction of the variance in the motion that can be explained by the OMN activity. If the fixational eye motion is generated by noise in the OMN activity, and the noise is independent in different OMNs, then the variance in the eye motion is a sum over contributions from individual OMNs. Intuitively, the sum over explained variances cannot exceed the actual variance. More formally, it is straightforward to see that for a set of independent random variables {*i*}, whose coefficients of determination with some other variable are denoted by 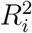, the sum 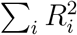 is bounded from above by unity (in our case, the independent random variables are the predictors of eye motion generated by each OMN, and the other variable is the actual motion of the eye). We conclude that if there are *N_m_* OMNs that independently drive eye motion,

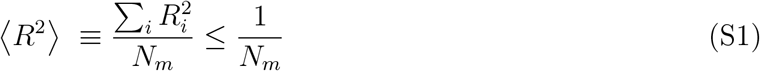

where 〈〉 denotes averaging over the population of OMNs.

Under the assumption that the recorded OMNs are representative of the full population, we can attempt to estimate 〈*R*^2^〉 based on our data. In practice, we found that it was difficult to obtain a reliable estimate of *R*^2^ directly from our data set, mostly due the large uncertaintly in the estimates of the covariances. We were able to obtain a more reliable estimate for the mean of 〈*R*〉, which could then be used to obtain a lower bound:

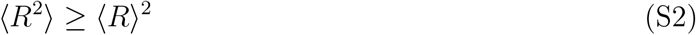

Since *N_m_* is estimated to be in the order of magnitude of a few thousands [1, 2], we would expect, based on Eqs. S1-S2 to obtain 〈*R*〉^2^ ≲ 10^−3^, which is an order of magnitude smaller than our estimates. Thus, we can reject the hypothesis that the fixational eye motion originates from independent contributions of noise in the individual OMNs.

### Contribution of oculomotor state to the CV of OMNs

We assessed the spiking variability of OMNs by examining the CV of the inter-spike interval during fixation. In doing so, we implicitly assumed that the firing rate remains fixed during each fixational epoch over which the CV was measured. However, we have argued in this work that the firing rate varies during the fixational epoch, due to the diffusive dynamics within the oculomotor integrator. This raises a question, whether the diffusive dynamics within the oculomotor integrator affects our measurements of the CV. Here we show that the effect of the diffusion on the measurement of the CV can be safely neglected.

Let us assume that the instantaneous firing rate of a neuron is given by

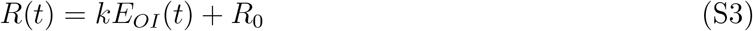

which is a random variable due to the variability of *E_OI_*, and that the CV is measured over an interval of duration Δ*t*.

To estimate the CV of the ISI that arises from the diffusion in the oculommotor integrator, let us assume that there is no intrinsic spiking variability: hence, the inter-spike interval is given precisely by 1/*R*. Assuming that the state of the integrator undergoes simple diffusion with diffusivity *D*, the variance of *E_OI_* over the measurement interval is then given by

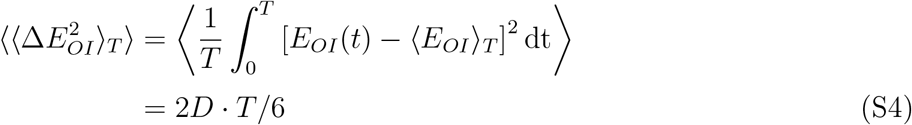

where averaging is with respect to time 〈…〉_*T*_, and over instances of the random diffusive trajectory of *E_OI_*. The CV of 1/*R* is then given by

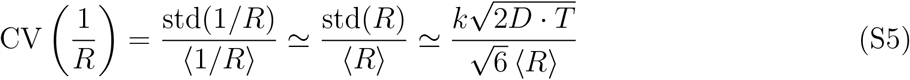

In a typical fixation interval of 500 ms, with *k* ~ 1/15 spikes/s/arcmin, 2D ≈ 100 arcmin^2^/s and 〈*R*〉 = 80 Hz, we conclude that the CV of the inter-spike interval is given by

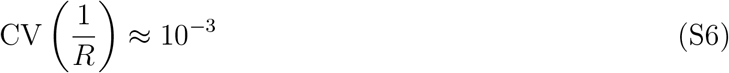

which is an order of magnitude smaller than the measured CV of OMNs. In addition note that under these biologically plausible assumptions 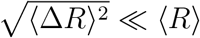. Hence, the approximations made in the above derivation are valid. We conclude that the contribution of the diffusive state of the integrator is negligible compared to other sources of noise in the OMNs. Similar argumentation holds for neurons within the oculomotor integrator.

### Contributions of peripheral noise and feedback to the eye motion

#### Influence of spiking noise on the MSD curve

Our model includes spiking noise in both the integrator neurons and OMNs. The primary role of spiking noise is to drive diffusive dynamics along the continuous attractor of the OI. A secondary effect of spiking noise within the OI is to generate a noisy output to OMNs, which exists even if the internal state of the OI is fixed (i.e., in the absence of diffusion). Here we demonstrate that this latter source of variability, and the intrinsic spiking variability of the OMNs both have a negligible effect on the MSD curve over the experimentally relevant time scales. To show this we performed simulations in which the spiking noise in OMNs and in the output of the oculomotor integrator were eliminated by (1) simulating the dynamics of the oculmotor integrator with spiking neurons to drive diffusion, but using the instantaneous firing rates of oculomtor neurons (as determined by their synaptic input) instead of spikes as an input to the OMNs, and (2) using deterministic OMNs. Results are shown in Extended Data Fig.10 where we compare the full model against the model without the spiking noise. Note that relative to other sources of motion the spiking noise is consequential only at very short time scales ~ 10^−3^ – 10^−2^ s. Beyond time lags of order 10^−2^ s the MSD curves with and without spiking noise are almost indistinguishable.

#### Visual feedback

As shown in the main text the diffusive dynamics within the oculomotor integrator, combined with the mechanic response properties of the ocular muscles and plant, is sufficient to explain the main qualitative features of the MSD curve: the superdiffusive nature of the motion up to time scales of at least several hundred ms, and the reduction in the logarithmic slope of the curve (to a value that still exceeds unity) beyond time scales that exceed approximately 100 ms. To obtain precise quantitative agreeement between the MSD curve predicted by the model and the experimetntally measured curve, we found that it was necessary to include a mechanism that decreases the slope of the MSD curve at time scales exceeding 100ms, even beyond the decrease in the slope that was already predicted based on the other ingredients of the model.

We thus included in the model a simple yet biologically plausible visual feedback mechanism that suppresses motion of the target over the retina. The features of this feedback mechanism were motivated as follows. First, several studies have shown that the statistics of eye motion during fixational drift are influenced by the visual stimulus [3, 4, 5, 6, 7, 8, 9]. This indicates that fixational drift is influenced, at least to some extent, by the stimulus. Second, recent measurements of fixational motion in human subjects using optical eye trackers were shown to exhibit weak oscillatory behavior over long time scales, with a period of about 100-200 ms [10, 11]. These oscillatory features could be explained by the existence of a delayed visual feedback mechanism [10]. The oscillations identified in [10] are reminiscent of similar oscillations that were previously observed during smooth pursuit [12]. These observations have led to our simple model of the visual feedback in which: (i) the brain generates an estimate of the recent drift velocity of the target on the retina by comparing a delayed estimate of the position of the target with the internal state of the OI, and (ii) this estimate, multiplied by a negative numerical prefactor, is fed as a velocity signal into the OI which drives motion in the opposite direction (Equations 27 and 17 in the main text).

#### Influence of the visual feedback on the MSD curve

The velocity feedback included in our model has several effects, illustrated in Extended Data Fig.4, which shows the MSD curve (Extended Data Fig.4a) and its logarithmic derivative (Extended Data Fig.4b) with and without visual feedback. For simplicity, and due to the very minor effect of OMN spiking noise on the MSD curves, we did not include spiking noise in these simulations. First, the visual feedback mechanism decreases the amplitude of the eye motion at all time lags (Extended Data Fig.4a). Second, it decreases the slope of the MSD curve (Extended Data Fig.4b). In terms of fitting the model to the data, the crucial effect of the feedback was the decrease in the slope of the MSD curve in comparison to the no-feedback case. This enables the model to fit well the MSD curve at both short and long time lags. Without the feedback we were only enable to fit well with the data in either short or long time lags (Extended Data Fig.5). In addition, the feedback decreases the amplitude of the MSD curve which facilitated the reach of a good match between the model and the data with biologically plausible choices for the parameters that control the magnitude of the diffusive motion, i.e., with 15K neurons in the integrator with CV of ISI of 0.20, and 1K OMNs.

